# Cell-specific expression of key mitochondrial enzymes precludes OXPHOS in astrocytes of the adult human neocortex and hippocampal formation

**DOI:** 10.1101/2022.10.27.514048

**Authors:** Arpad Dobolyi, Attila G Bago, Anna Stepanova, Krisztina Paal, Jeonghyoun Lee, Miklos Palkovits, Christos Chinopoulos

## Abstract

The astrocyte-to-neuron lactate shuttle hypothesis entails that glycolytically derived pyruvate in astrocytes is converted to lactate instead of being catabolized in mitochondria. The mechanism of this metabolic rewiring is unclear. Here we show that astrocytes of the adult human neocortex and hippocampal formation do not express mitochondrial proteins critical for performing oxidative phosphorylation (OXPHOS) to a detectable degree, including cytochrome c and complex IV. Without OXPHOS, human brain astrocytes are bound to produce lactate to avoid interruption of glycolysis.

## Main

Simply stated, the Astrocyte-To-Neuron Lactate Shuttle (ANLS) model dictates that in response to glutamate-mediated neuronal activity, astrocytes enhance their glycolytic flux forming lactate, which is shuttled to neurons through monocarboxylate transporters ^1^. An understated but key-point of this model is that in astrocytes, glycolytically produced pyruvate is converted to lactate instead of entering mitochondria and get oxidatively catabolized, even in the presence of oxygen ^2^. This evolutionary conserved ^3^ astrocyte-specific “aerobic glycolysis”-more commonly associated with cancer ^4^-has never been adequately explained. In the original scheme of the article proposing the ANLS hypothesis, mitochondria were tactfully missing from astrocytes ^1^, and this has remained unchanged ever since ^5^, ^6^. Pellerin and Magistretti attributed the conversion of pyruvate into lactate in astrocytes to a lack of mitochondrial aspartate/glutamate carrier that reduces their capacity to transfer NADH by the malate/aspartate shuttle (MAS) in the mitochondria in order to regenerate NAD^+5^. Thus, lactate dehydrogenase would be essential for regenerating NAD^+^ which is indispensable for glyceraldehyde dehydrogenase, allowing glycolysis to proceed ^7^.

However, the absence of MAS does not universally imply upregulation of glycolysis, see for example ^8^; furthermore, MAS inhibition leads only to an incomplete prevention of mitochondrial utilization of pyruvate ^9^. Finally, it was recently discovered that cytosolic malate dehydrogenase (MDH1) may also assume the role of regenerating NAD^+^ when mitochondria are dysfunctional ^10^. The increase in astrocytic glycolytic flux could be explained via a mechanism involving Na^+^/K^+^ ATPase ^1^, however that could not account for pyruvate not getting metabolized in mitochondria. The groups of Bolaños, Felipe Barros and Clark have offered the explanation that NO-which is formed and released by neurons during glutamatergic neurotransmission ^11^-activates glycolysis ^12^ simultaneously to inhibiting complex IV in astrocytes ^13^ at nanomolar concentrations ^14^ by reversibly competing with oxygen^15^. These effects cause the activation of glycolysis in astrocytes releasing lactate ^16^. The groups of Bolaños have also reported that murine neuronal mitochondrial complex I is assembled into supercomplexes increasing OXPHOS efficacy, whereas the majority of astrocytic complex I is free ^17^; in view of this astrocytic OXPHOS inefficiency, there should be pyruvate-> lactate formation in order to regenerate cytosolic NAD^+^. Finally, the group of Sokoloff published that the activity of rat astrocytic pyruvate dehydrogenase complex is low due to hyperphosphorylation, unfavoring mitochondrial utilization of pyruvate, thus diverting it towards lactate ^18^.

The consensus that astrocytic energy demands are mostly met by glycolysis, whereas those of neurons mainly rely on OXPHOS is almost universally accepted (see ^16^ for in-favour and ^19^ for against); however, it is entirely based on work from cells or brain tissues of laboratory rodents ^1^, ^20^, ^21^. On the other hand, the group of McKenna demonstrated that murine astrocytes exhibit the capacity for oxidative decarboxylation of glutamate ^22^, which is consistent with the finding that mouse astrocytes express components of the citric acid cycle and the electron transport chain *in vivo* ^23^,^24^. Despite that the ANLS model has been demonstrated in human brain *in vivo* ^25^, Dienel and Yellen have been questioning its validity ^26^, ^27^, mostly on the grounds that murine astrocytes exhibit the capacity for performing the citric acid cycle and oxidative phosphorylation on the basis of transcriptomic, proteomic and functional analysis. However, the group of Chinopoulos showed that subunits coding for succinyl-CoA ligase and the α-ketoglutarate dehydrogenase complex (KGDHC)-two entities of the citric acid cycle-are lacking from adult human brain astrocytes ^28^, ^29^. Lack of KGDHC immunoreactivity from human glia has also been reported earlier by Blass ^30^. This severely truncated citric acid cycle present in adult human astrocytes, in conjunction to the reports that astrocytic glycolysis yields lactate with little-if any-entry into mitochondria, led us to investigate the cell-specific expression of OXPHOS-critical enzymes.

All antibodies used in this study have been firmly established as valid, monospecific antibodies for the techniques and tissue types used hereby. Proteins required for OXPHOS, cytochrome c and Cox IV were found only in the neurons based on the large size of the labelled cell bodies and also the morphology of the cell bodies and visible primary dendrites. These characteristics of cytochrome c and Cox IV protein expression were the same in the subiculum (Fig. 1), the dentate gyrus (Suppl. Fig. 1), and the parahippocampal cortex (Suppl. Fig. 2). The presence of labelled cell bodies in grey but not white matter confirmed the neuronal but not astrocytic localization in human brain. Double immunolabelling with the astrocytic marker S100 protein also demonstrated that astrocytes do not contain cytochrome c and Cox IV proteins to a detectable degree (Fig. 1F). Our data are entirely congruent with those reporting Cox IV cell-specific enzymatic activity or immunoreactivity *in situ* in animals placed in high positions in the phylogenetic tree; furthermore, the higher the position of the organism in the phylogenetic tree, the less the Cox IV enzymatic activity reported: specifically, in auditory relay nuclei of cat, Cox IV activity was almost exclusively (but not completely) expressed in neurons, than in non-neuronal neuropil ^31^; in the Lateral Geniculate Nucleus of cat, Cox IV immunoreactivity was present only in the neurons ^32^; In monkey hippocampus, Cox IV immunoreactivity was not present in glia ^33^ (in this reference, the same was also reported for mouse cerebellum); in macaque cerebellum and hippocampus, very low Cox IV reactivity was reported in non-neuronally populated areas ^34^; in primate visual cortex of macaque <2% of Cox IV immunoreactivity was present in glia in ^35^; in the lateral entorhinal cortex of the rhesus monkey, Cox IV immunoreactivity was detected only in the neurons; In human entorhinal cortex, Cox IV immunoreactivity was not present in glial elements ^36^; In ^37^, no Cox IV enzymatic activity staining was observed in the developing and adult human cortical layers where non-neuronal elements are found. In ^38^ it was shown that Cox IV mRNA was localized mainly or exclusively in cell bodies and proximal dendrites; this is consistent with our investigation, showing that mRNA levels obtained from the Allen Institute for Brain Science suggested that the mRNAs of cytochrome c and Cox IV are expressed in neurons to a much higher extent than in non-neuronal cell types (Fig. 1A). In contrast to the absence of immunolabelling of astrocytic Cox IV in the human cortical areas, Cox IV was found in both neuronal and glial cell types in the rat cerebral cortex (Suppl. Fig. 3). Heatmaps of mRNA expression levels in neuronal (Neu) and non-neuronal (N-N) cells based on single cell sequencing by the Allen Institute for Brain Science for all subunits participating in OXPHOS in humans (*homo*) vs mice (*mus*) is depicted in supplementary datasets 4 and 5, respectively.

**Figure 1.**
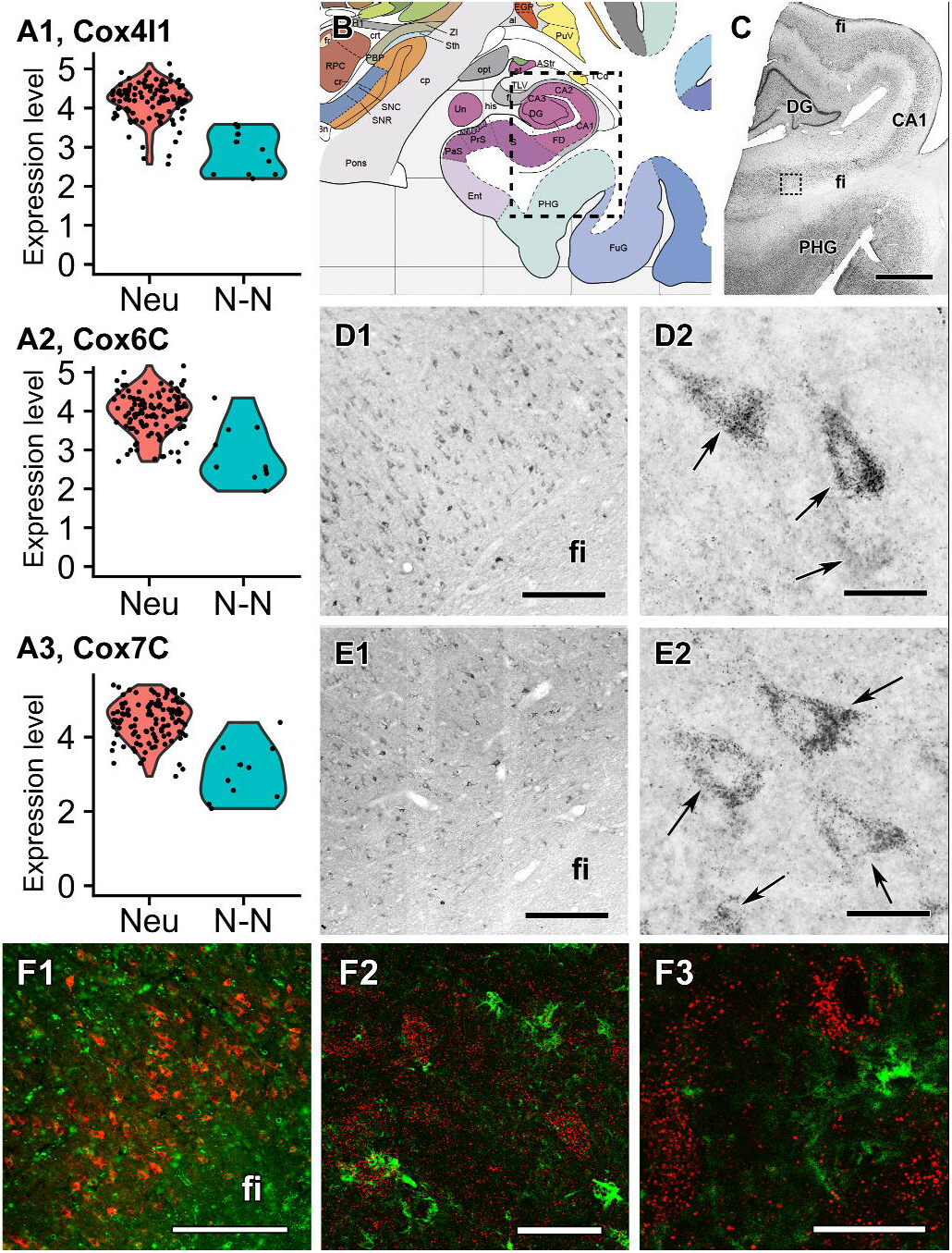
The expression of CoxIV and cytochrome c in neuronal cells of the subiculum. A: Violin plots of expression of mRNA level in neuronal (Neu) and non-neuronal (N-N) cells based on single cell sequencing of Allen Institute for Brain Science. The dots represent individual cells. A1-3 panel demonstrate a dominantly neuronal expression of different CoxIV subunits. B: Drawing of a coronal plane from a human brain atlas ^48^. C: A Nissl stained section corresponding to the area framed in panel A. D: Parts of a CoxIV-immunolabelled coronal section, which is parallel to the section shown in panel B. D1 shows the subicular area framed in panel C while D2 shows CoxIV-immunolabelled neurons at high magnification. E: Parts of a Cytochrome c immunolabelled coronal section, which is parallel to the section shown in panel C. E1 shows the area framed in panel C while E2 shows Cytochrome c immunolabelled cells at high magnification. In D2 and E2, arrows point to individual labelled cells, which are neurons based on their size and morphology. Also note the absence of labelling of cells with glial morphology. Also note the dot-like distribution of labelling suggesting the location of the enzymes in mitochondria. In addition, note the absence of labelled cells in the white matter of fimbria hippocampi (fi). F: Verification of non-glial location of CoxIV (red) using double immunolabelling with the astrocyte marker S100 (green). Low magnification image (F1) shows that only S100 but not CoxIV is present in the fimbria hippocampi (fi). A higher magnification image (F2) demonstrates the lack of co-localization between CoxIV and S100. The high magnification image shows the dot-like mitochondrial distribution of CoxIV. Further abbreviations: CA1 = CA1 region of the hippocampus, DG = dentate gyrus, PHG = parahippocampal gyrus. Scale bars = 3 mm for C, 300 μm for D1-F1, 50 μm for F2, 30 μm for D2 and E2, and 20 μm for F3.

As opposed to cytochrome c and Cox IV, pyruvate carboxylase was found exclusively in astrocytes based on the morphology and localization of the cells (Fig. 2A) as well as double labelling with S100 (Fig. 2G) consistent with literature on mice ^39^. Pyruvate transporters were found in both neuronal and glial cell types, but with differential expression as MPC1 was found only in neurons while MPC2 only in astrocytes in the subiculum (Fig. 2B and C), the dentate gyrus (Suppl. Fig. 1) and the parahippocampal gyrus (Suppl. Fig. 2). While this pattern of expression is at odds with ^40^, we confirmed the exclusively neuronal localization of MCP1 using double labelling with S100 as well (Fig. 2H). Additional pyruvate utilization enzymes are present in both neurons and astrocytes as we found 3 different subunits of pyruvate dehydrogenase (PDH) localized in both cell types in the subiculum (Fig. 2D-F), the dentate gyrus (Suppl. Fig. 1) and the parahippocampal cortex, too (Suppl. Fig. 2).

**Figure 2.**
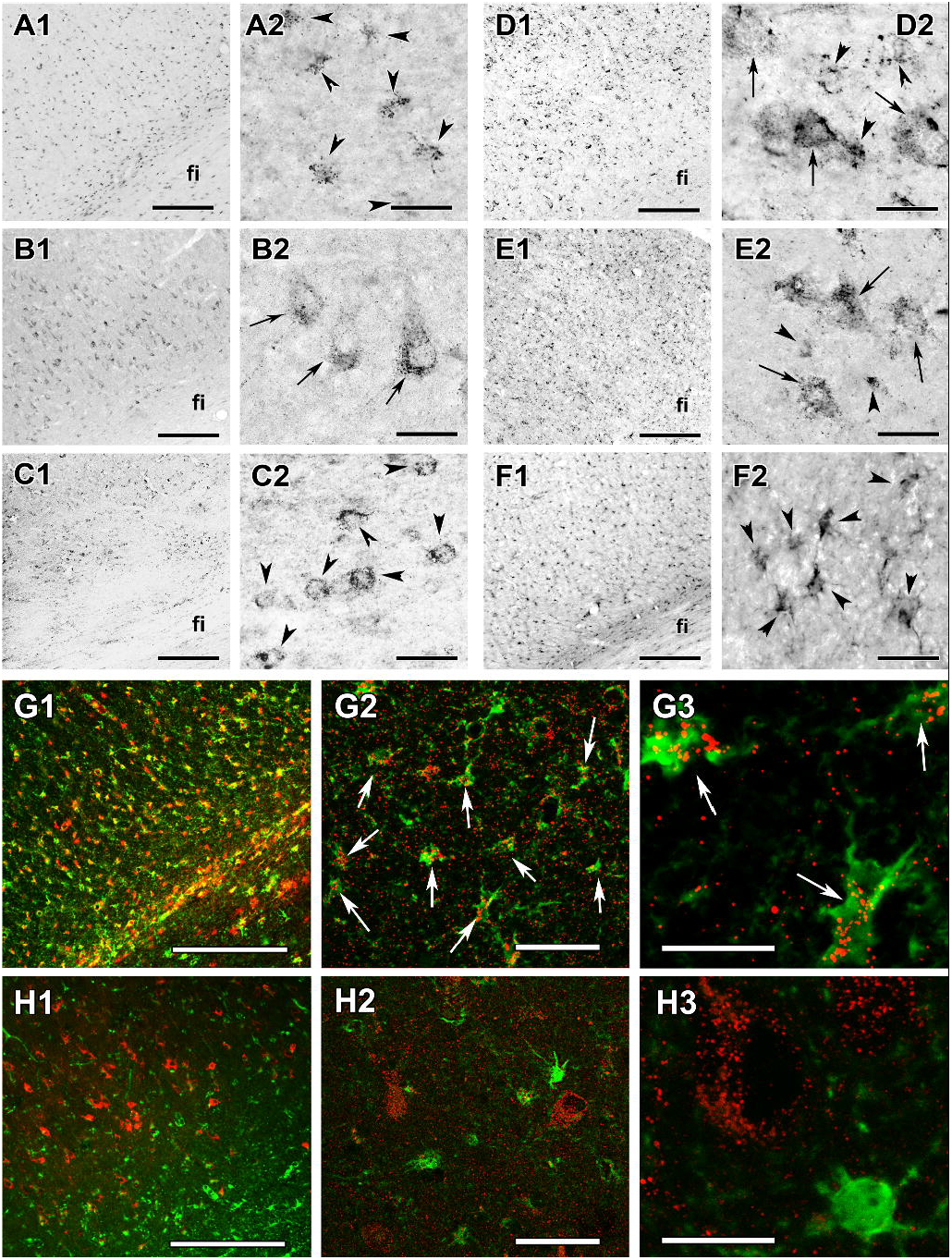
Immunolabelling of pyruvate carboxylase, MPC1, MPC2, and PDH enzymes in the subiculum. The same brain areas are shown as in figure 1. Left panels indicate pictures taken at low while right panels next to them taken at high magnification. In the high magnification images, arrows point to neurons while arrowheads point to astrocytes. A: Parts of a pyruvate carboxylase immunolabelled coronal section. Arrowheads in A2 point to individual labelled cells, which resemble astrocytes based on the relatively small size of their cell bodies as well as their morphology. Note the dot-like distribution of labelling consistent with the mitochondrial location of the enzymes. B: MPC1-immunolabelled sections show labelled neurons based on their exclusive distribution in the grey matter (see the absence of labelled cells in the white matter of fimbria hippocampi (fi) in B1 as opposed to their presence in A1), and also on their size and morphology (B2). C: MPC2-immunolabelled section. Labelled cells are present in the white matter of the fi (C1). High magnification picture (C2) shows relatively small labelled cells, likely astrocytes. D: PDHE1b, E: PDHE2, F: PDHX. Low magnification pictures on the left (D1, E1, F1) as well as high magnification figures on the right show that PDHE1b and PDHE2 are expressed in both neurons and glial cells based on the morphology of labelled cells while PDHX expression is exclusively glial. G: Demonstration of glial location of pyruvate carboxylase (red) using double immunolabelling with the astrocyte marker S100 (green). Relatively low magnification image (G1) shows that S100 and pyruvate carboxylase are present in the grey as well as in the white matter (fimbria hippocampi - fi). The higher magnification image (G2) demonstrates that all cells are double labelled with pyruvate carboxylase and S100 (pointed to by white arrows). The high magnification image in the right panel (G3) shows the dot-like mitochondrial distributions of pyruvate carboxylase. Astrocytes visualized by S100 labelling (green) contain pyruvate carboxylase-containing mitochondria (red). H: Demonstration of non-glial location of MPC1 (red) using double immunolabelling with the astrocyte marker S100 (green). The low magnification image in the left panel (H1) shows that S100-immunolabelled are present in the white matter of fi while MPC1-immunoreactivity is absent in this region. The higher magnification images demonstrate the lack of double labelling between MPC1 and S100 (H2, H3). Scale bars = 300 μm for A1-H1, 50 μm for G2 and H2, 30 μm for A2-F2, and 20 μm for G3 and H3.

The MDH2 of the tricarboxylic acid cycle was also not present in human astrocytes but only in neurons in the subiculum (Fig. 3A), the dentate gyrus (Suppl. Fig. 1) and the parahippocampal cortex (Suppl. Fig. 2). This, together with the fact that human astrocytes do not express KGDHC ^29^ essentially mitigates any reasoning for having OXPHOS in astrocytes. Therefore, it is safe to assume that O_2_ consumption in human brain should be attributed to only neuronal mitochondria; mindful that in the human brain the glia-to-neuron ratio is close to one ^41^, it can be stated that 50% of the human brain is responsible for the 20 % of the total body oxygen consumption ^42^, thus, only 1 % of body weight. With the current state-of-the-art technological advances regarding *in vivo* functional assessment, it is not possible to verify that only neurons contribute to O_2_ consumption in the living human brain; in the most recent, most advanced MRI of the human brain exhibiting a spatial resolution of 200 μm, 253-264 neuronal and glial bodies are included ^43^. Having said that, it is notable that non-oxidative consumption of glucose during focal physiologic neural activity has been reported in 1988 ^44^, i.e. six years before ANLS hypothesis was formulated ^1^, that has led to the establishment of the concept of regional aerobic glycolysis in the human brain ^45^.

**Figure 3.**
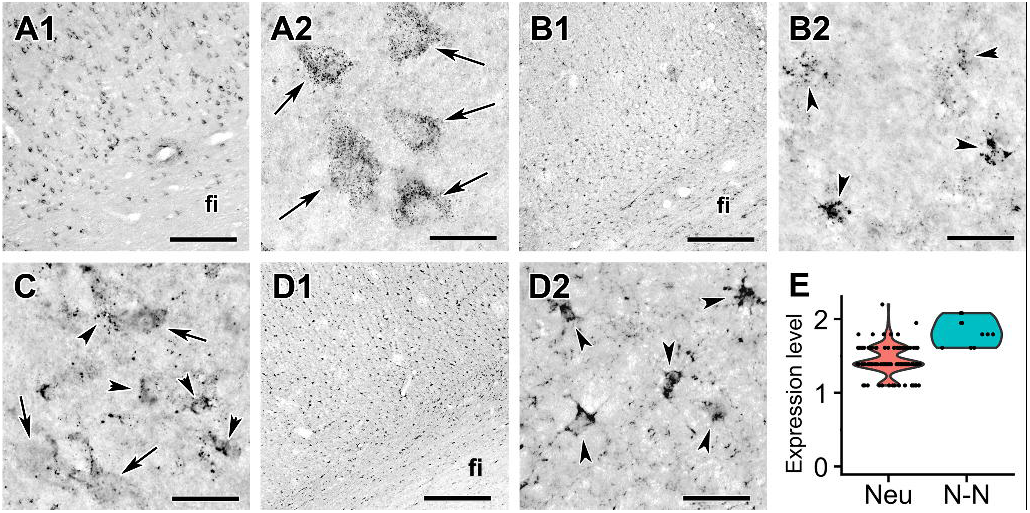
Labelling of mitochondrial dehydrogenases in the subiculum. Arrows point to neurons while arrowheads indicate glial cells. Note the dot-like mitochondrial labelling. A: Low magnification picture on the left (A1) as well as high magnification figure on the right (A2) show that MDH2 is expressed exclusively in neurons. B: Low magnification image on the left (B2) as well as high magnification image on the right show that IDH2 expression is exclusively glial. C: A high magnification image demonstrated that SDH1-B is expressed in both neurons and glial cells based on the morphology of labelled cells. D: SDH-C expression is exclusively glial. E: Violin plot of expression of mRNA level in neuronal (Neu) and non-neuronal (N-N) cells based on single cell sequencing of Allen Institute for Brain Science. The dots represent individual cells demonstrating a dominantly non-neuronal expression of SDH-C. Scale bars = 300 μm for A1, B1 and D1, and 30 μm for A2, B2, C and D2.

The lack of Cox IV from adult human glia should not be too surprising to the scientific community using rodents as laboratory animals: mice engineered to lack Cox IV in astrocytic mitochondria *in vivo* were fully viable in the absence of any signs of glial or neuronal loss even at 1 year of age ^46^.

In turn, we found that SDH1B was present in both neuronal and glial cell types in the subiculum (Fig. 3B), the dentate gyrus (Suppl. Fig. 1) and also in the parahippocampal cortex (Suppl. Fig. 2), which is expected, given the multiple roles of succinate in non-OXPHOS pathways ^47^.

## Methods

### Human brain tissue samples

Human brain samples were collected in the Human Brain Tissue Bank, Semmelweis University in accordance with the Ethical Rules for Using Human Tissues for Medical Research in Hungary (HM 34/1999) and the Code of Ethics of the World Medical Association (Declaration of Helsinki). Tissue samples were taken during brain autopsy at the Department of Forensic Medicine of Semmelweis University in the framework of the Human Brain Tissue Bank (HBTB), Budapest. The activity of the HBTB has been authorized by the Committee of Science and Research Ethic of the Ministry of Health Hungary (ETT TUKEB: 189/KO/02.6008/2002/ETT) and the Semmelweis University Regional Committee of Science and Research Ethic (No. 32/1992/TUKEB). The study reported in the manuscript was performed according to a protocol approved by the Committee of Science and Research Ethics, Semmelweis University (TUKEB 189/2015). The medical history of the subjects was obtained from clinical records, interviews with family members and relatives, as well as pathological and neuropathological reports. All personal data are stored in strict ethical control, and samples were coded before the analyses of tissue.

### Tissue collection for immunolabelling

Immunohistochemistry was used to assess the location of mitochondrial proteins in the hippocampus and the adjacent portion of the parahippocampal gyrus. For immunolabelling, brain blocks from a 62 years old female and a 56 years old male individual was cut into 10 mm thick coronal slice and immersion fixed in 4% paraformaldehyde in 0.1 M phosphate-buffered saline (PBS) for 5 days. Subsequently, the block was transferred to PBS containing 0.1% sodium azide for 2 days to remove excess paraformaldehyde. Then, the block was placed in PBS containing 20% sucrose for 2 days of cryoprotection. The block was frozen and cut into 60 μm thick serial coronal sections on a sliding microtome. Sections were collected in PBS containing 0.1% sodium azide and stored at 4 °C until further processing.

### DAB immunolabelling

Free-floating brain sections were immunolabeled for mitochondrial proteins (see antibodies and their dilutions in Supplementary table 1). The antibodies were applied for 24 h at room temperature, followed by incubation of the sections in biotinylated anti-rabbit/mouse secondary antibody (1:1,000 dilution, Vector Laboratories, Burlingame, CA) and then in avidin-biotin-peroxidase complex (1:500, Vector Laboratories) for 2 h. Subsequently, the labelling was visualized by incubation in 0.02% 3,3-diaminobenzidine (DAB; Sigma), 0.08% nickel (II) sulphate and 0.001% hydrogen peroxide in PBS, pH 7.4 for 5 min. Sections were mounted, dehydrated and coverslipped with Cytoseal 60 (Stephens Scientific, Riverdale, NJ, USA).

### Double labelling with the glial marker S100

Double immunofluorescence staining was used to clarify the colocalization of mitochondrial genes with the glial marker S100 in the hippocampus and parahippocampal gyrus. To reduce autofluorescence, tissue sections were treated with 0.15% Sudan Black B (in 70% ethanol) after antigen retrieval (10 min at 90°C in 0.05 M Tris buffer, pH = 9.0) procedures. Slides were blocked by incubation in 3% bovine serum albumin (with 0.5% Triton X-100 dissolved in 0.1 M PB, Sigma) for 1 h at room temperature, followed by washing with washing buffer (10 min × 3). Mitochondrial proteins were immunolabeled for single labelling except for the visualization, which was performed with fluorescein isothiocyanate (FITC)-tyramide (1:8,000 dilution) and H2O_2_ (0.003%) in 100 mM Trizma buffer (pH 8.0 adjusted with HCl) for 6 min. Subsequently, sections were placed in mouse anti-S100 (1:250 dilution, Millipore, Cat. No. MAB079-1) for 24 h at room temperature. The sections were then incubated in Alexa 594 donkey anti-mouse secondary antibody (1:500 dilution, Vector Laboratories) followed by 2 h incubation in a solution containing avidin-biotin-peroxidase complex (ABC, 1:300 dilution, Vector Laboratories). Finally, all sections with fluorescent labels were mounted on positively charged slides (Superfrost Plus, Fisher Scientific, Pittsburgh, PA) and coverslipped in antifade medium (Prolong Antifade Kit, Molecular Probes).

### Microscopy and photography

Sections were examined using an Olympus BX60 light microscope also equipped with fluorescent epi- illumination and a dark-field condensor. Images were captured at 2048 × 2048 pixel resolution with a SPOT Xplorer digital CCD camera (Diagnostic Instruments, Sterling Heights, MI, USA) using a 4× objective for dark-field images, and 4–40× objectives for bright-field and fluorescent images. Confocal images were acquired with a Zeiss LSM 70 Confocal Microscope System using a 40-63X objectives at an optical thickness of 1 μm for counting varicosities and 3 μm for counting labelled cell bodies. Contrast and sharpness of the images were adjusted using the ‘levels’ and ‘sharpness’ commands in Adobe Photoshop CS 8.0. Full resolution was maintained until the photomicrographs were cropped at which point the images were adjusted to a resolution of at least 300 dpi.

## Supporting information

Dataset 4 (homo)

Dataset 5 (mus)

## Acknowledgements

This work was supported by grants from NKFIH KH129567, NKFIH K135027 and TKP2021-EGA-25 to C.C, NKP-2017-00002 to M.P., and NKFIH K134221, NAP2022-I-3/2022 to A.D.

