## Supplementary material for "Cell-specific expression of key mitochondrial enzymes precludes OXPHOS in astrocytes of the adult human neocortex and hippocampal formation": Legends to supplementary figures

**Supplementary figure 1.** *Labelling of mitochondrial enzymes in the dentate gyrus*. A: MPC1, B: MPC2, C: PDHE1B, D: PDHE2, E: PDHX, F: pyruvate carboxylase, G: IDH2, H: SDH1B, I: SDH-C, J: MDH2, K: Cytochrome C, L: CoxIV. MPC1, MDH2, Cytochrome c and CoxIV, immunostaining labels exclusively neurons, while MPC2, PDHX, pyruvate carboxylase, IDH2 and SDH-C label only astrocytes. In turn, PDHE1B, PDHE2 and SDH1B immunoreactivity is present in both neuronal and glial cells. The magnification of all panels is the equal. Scale bar: 100 µm.

**Supplementary figure 2.** *Labelling of selected mitochondrial enzymes in the parahippocampal gyrus*. A: CoxIV, B: Cytochrome C, C: MPC1, D: MDH2, E: pyruvate carboxylase, F: MPC2, F: PDHE1B, G: SDH1B. CoxIV, Cytochrome C, MPC1 and MDH2 immunostaining labels exclusively neurons (A-D), while pyruvate carboxylase and MPC2 labels only astrocytes. In turn, PDHE1B and SDH1B immunoreactivity is present in both neuronal and glial cells. The magnification of all panels is the equal. Scale bar: 100 µm.

**Supplementary figure 3.** *Neuronal and glial expression of CoxIV in the rat cerebral cortex.* A: DAB immunolabelled image shows several labelled neurons. In addition, potential astrocytes, pointed to by black arrows, are also labelled. B: Double labelling of CoxIV and the astrocyte marker S100 shows that in addition to intensely labelled neurons (green), S100 labelled astrocytes (red) also contain a moderate amount of CoxIV. The area indicated by the white arrow is enlarged in the inlet in the top right corner. Scale bars = 30 µm for A and 60 µm for B.

**Supplementary dataset 4.** *Heatmaps of mRNA expression levels in neuronal (Neu) and non-neuronal (N-N) cells based on single cell sequencing by the Allen Institute for Brain Science for all subunits participating in OXPHOS in human brain (homo)*. Queries were made by inputting gene symbols enlisted in MitoCarta 2.0 and curating it to include new data reported in MitoCarta 3.0. Heatmap range is identical for all panels as shown in the “homo Pi carriers and dicarboxylate carriers”.

**Supplementary dataset 5.** *Heatmaps of mRNA expression levels in neuronal (Neu) and non-neuronal (N-N) cells based on single cell sequencing by the Allen Institute for Brain Science for all subunits participating in OXPHOS in mouse brain (mus)*. Queries were made by inputting gene symbols enlisted in MitoCarta 2.0 and curating it to include new data reported in MitoCarta 3.0. Heatmap range is identical for all panels as shown in the “mus Pi carriers and dicarboxylate carriers”.
