## Supplementary figures and images for "Cell-specific expression of key mitochondrial enzymes precludes OXPHOS in astrocytes of the adult human neocortex and hippocampal formation"

### homo ANT.png

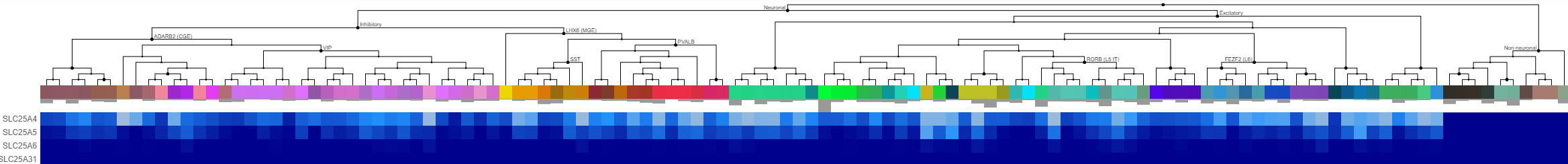

### homo complex I.png

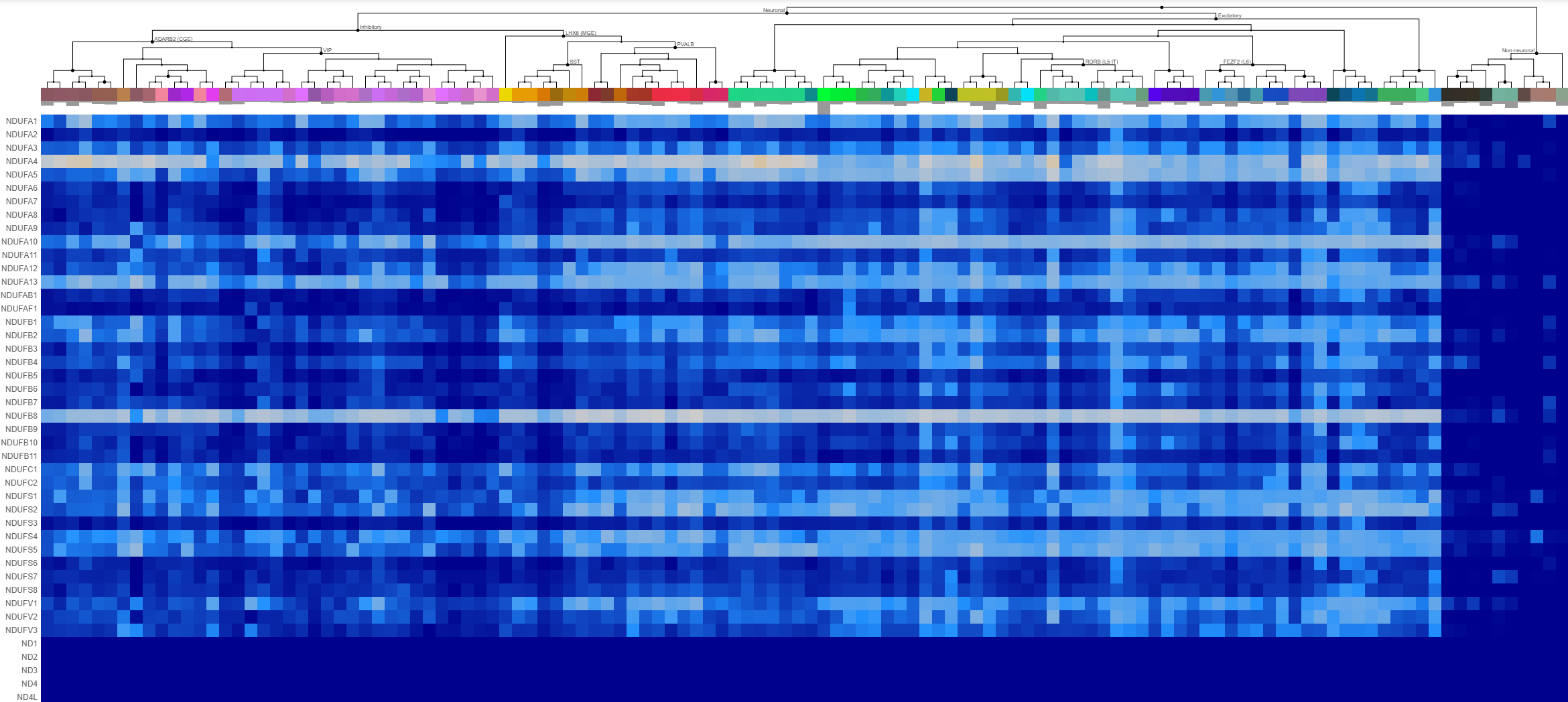

### homo complex I_remaining genes.png

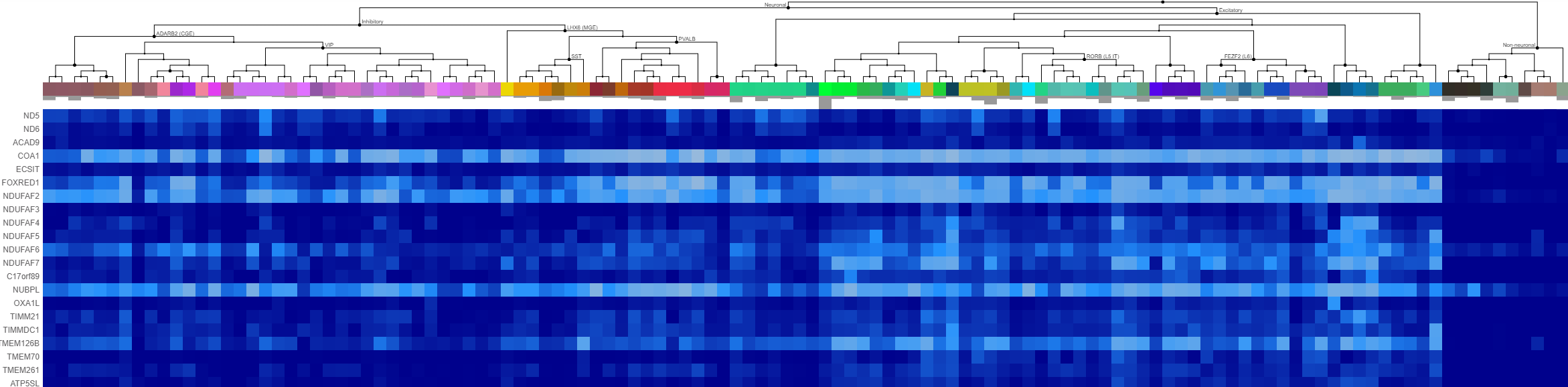

### homo complex II.png

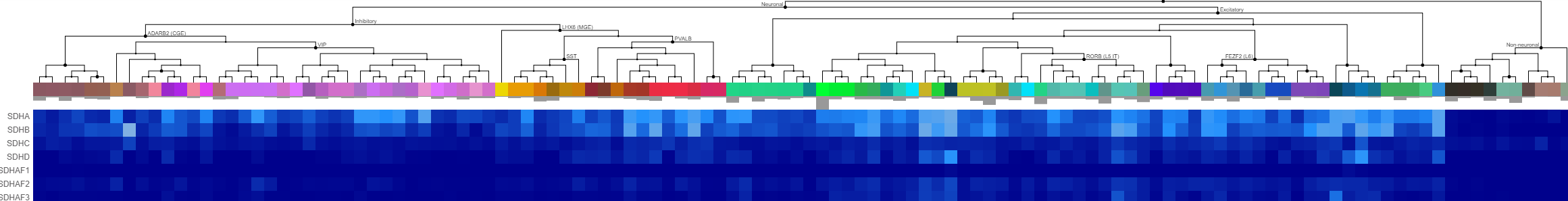

### homo complex III.png

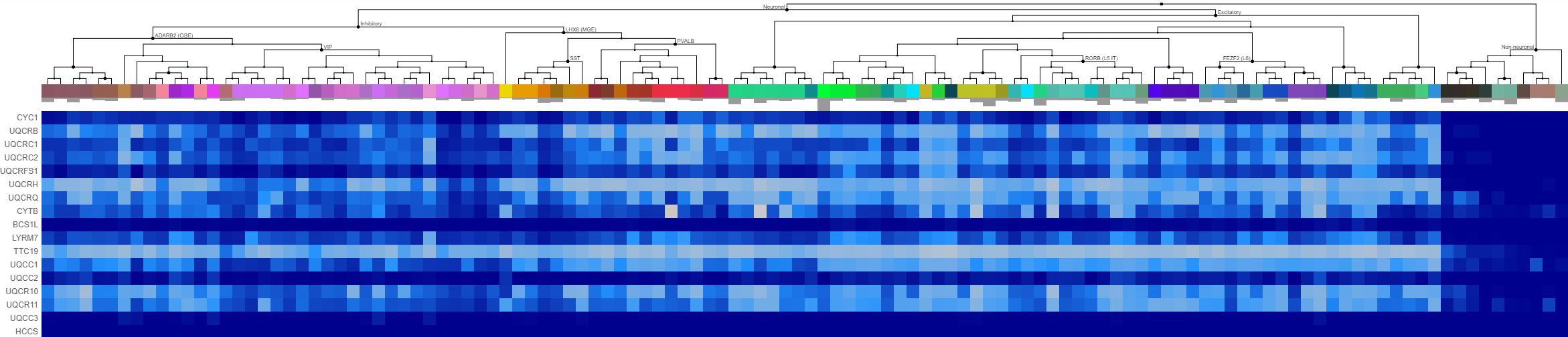

### homo complex IV pseudogenes.png

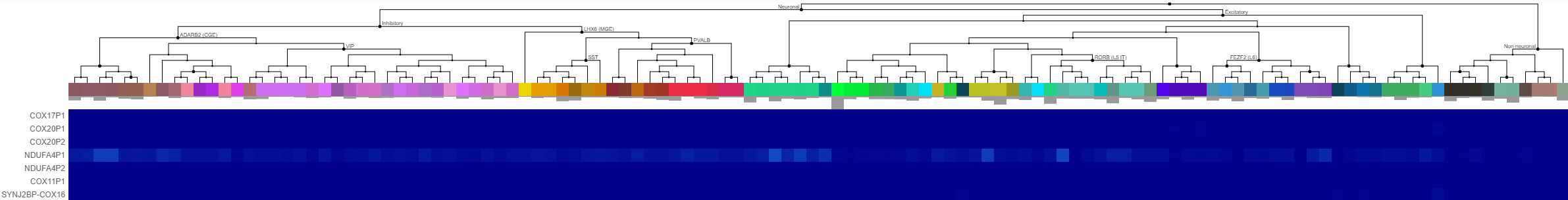

### homo complex IV.png

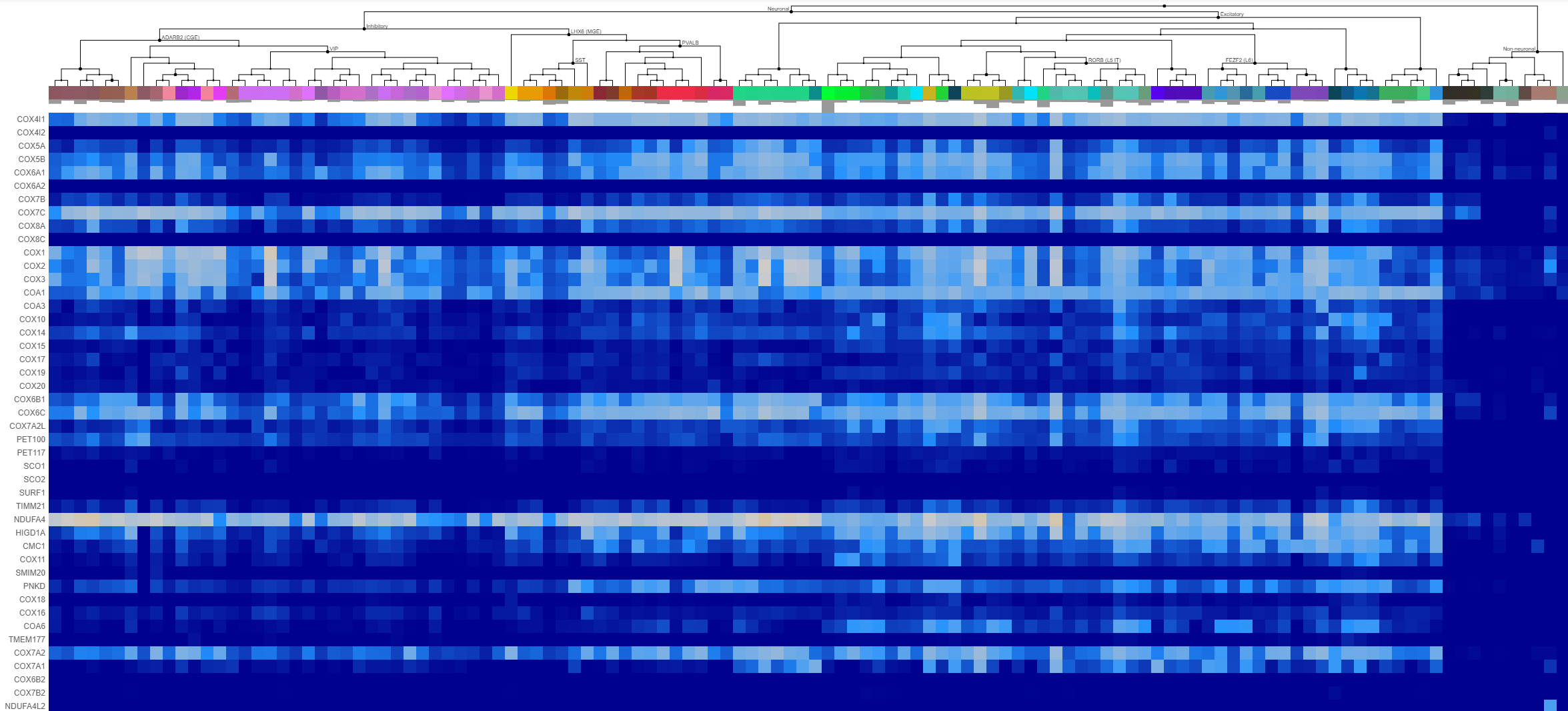

### homo complex V.png

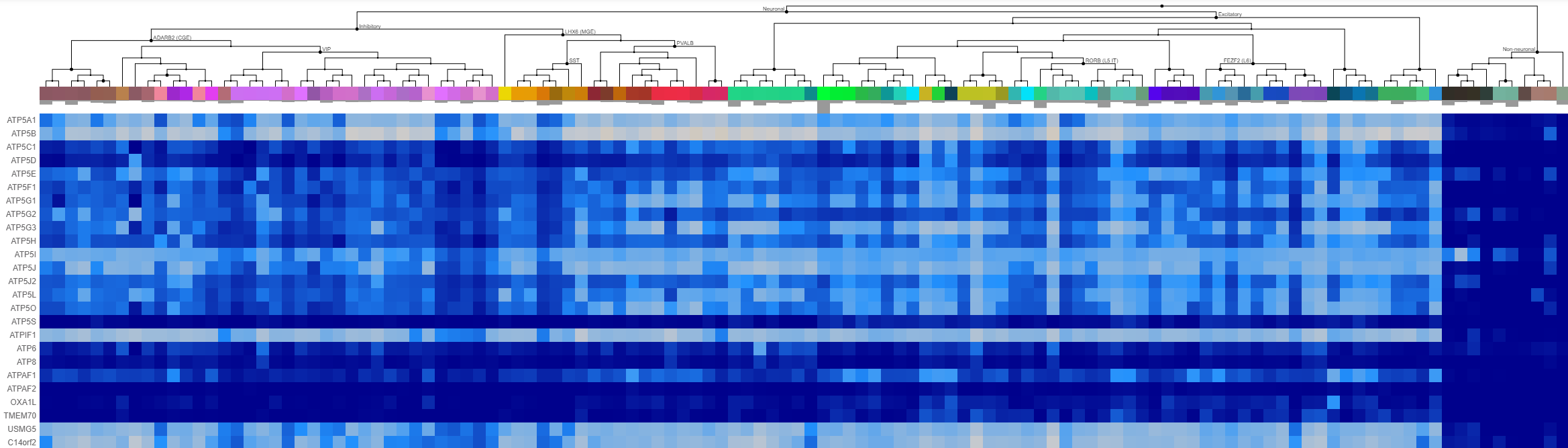

### homo cyt c.png

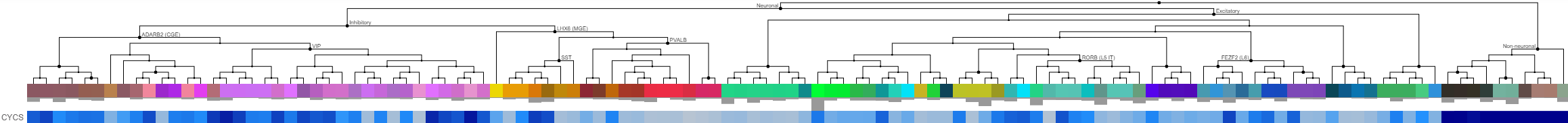

### homo MDH2 and IDH.png

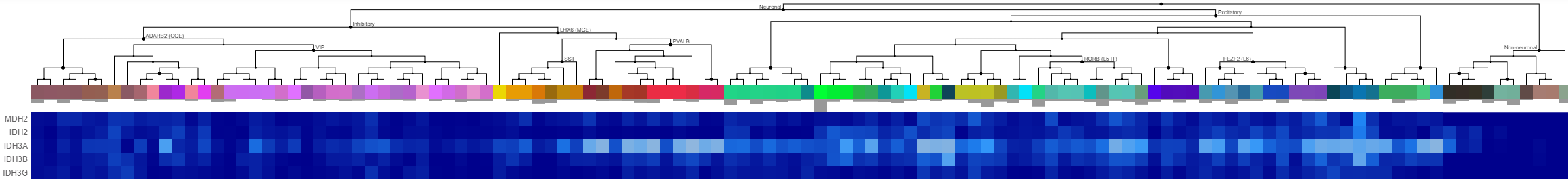

### homo PDH.png

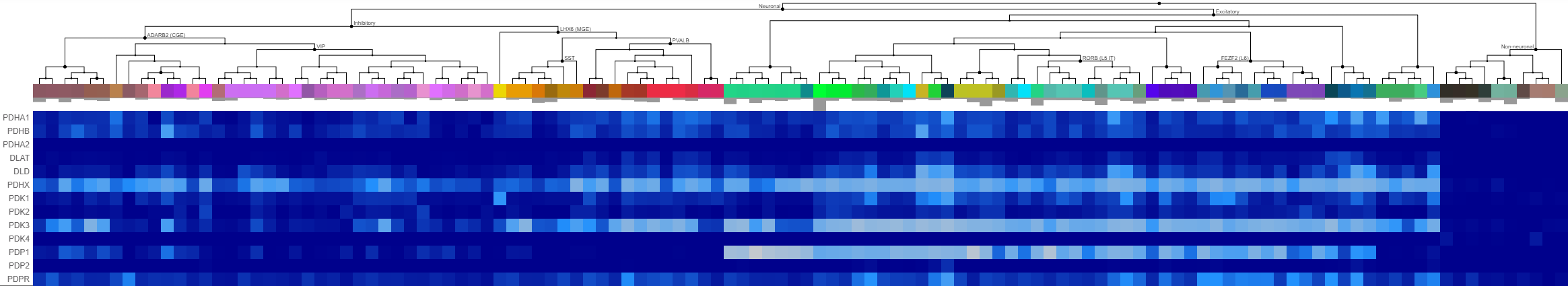

### homo Pi carriers and dicarboxylate carriers.png

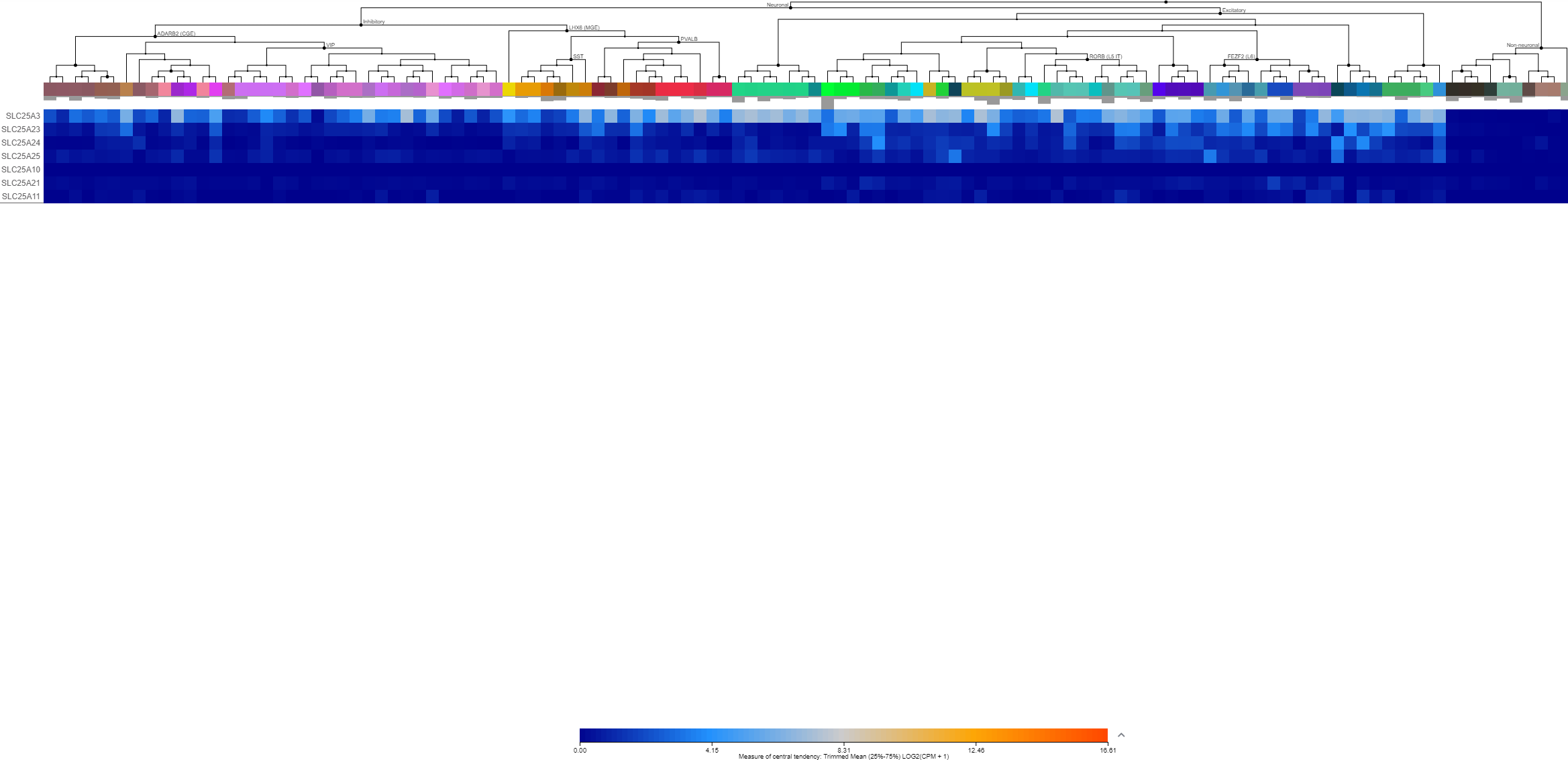

### mus ANT.png

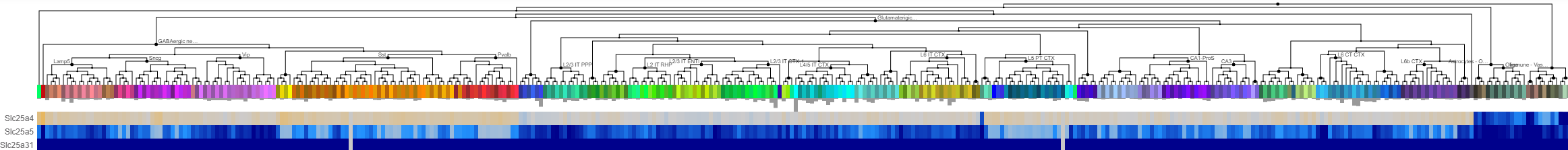

### mus complex I remaining genes.png

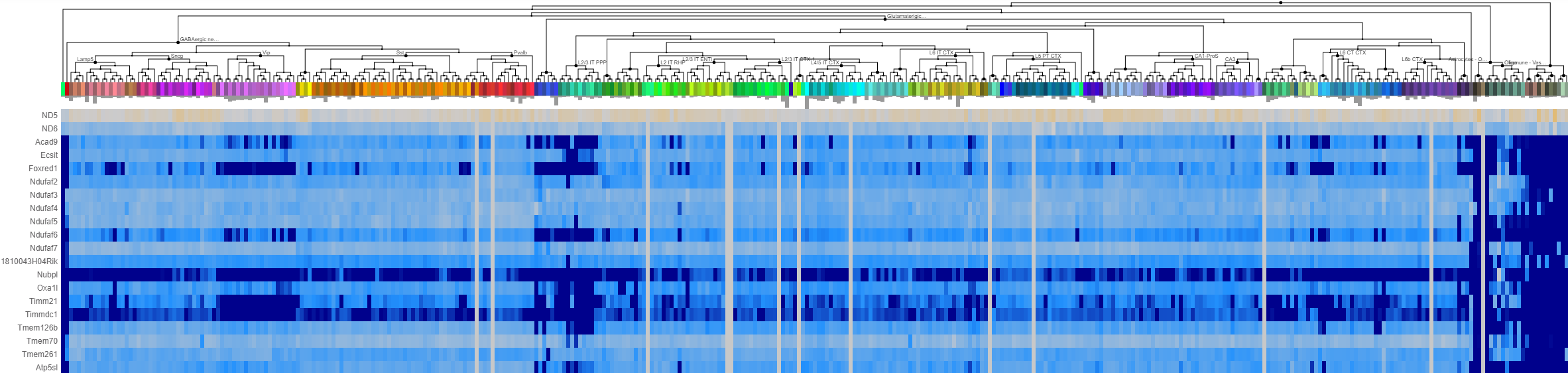

### mus complex I.png

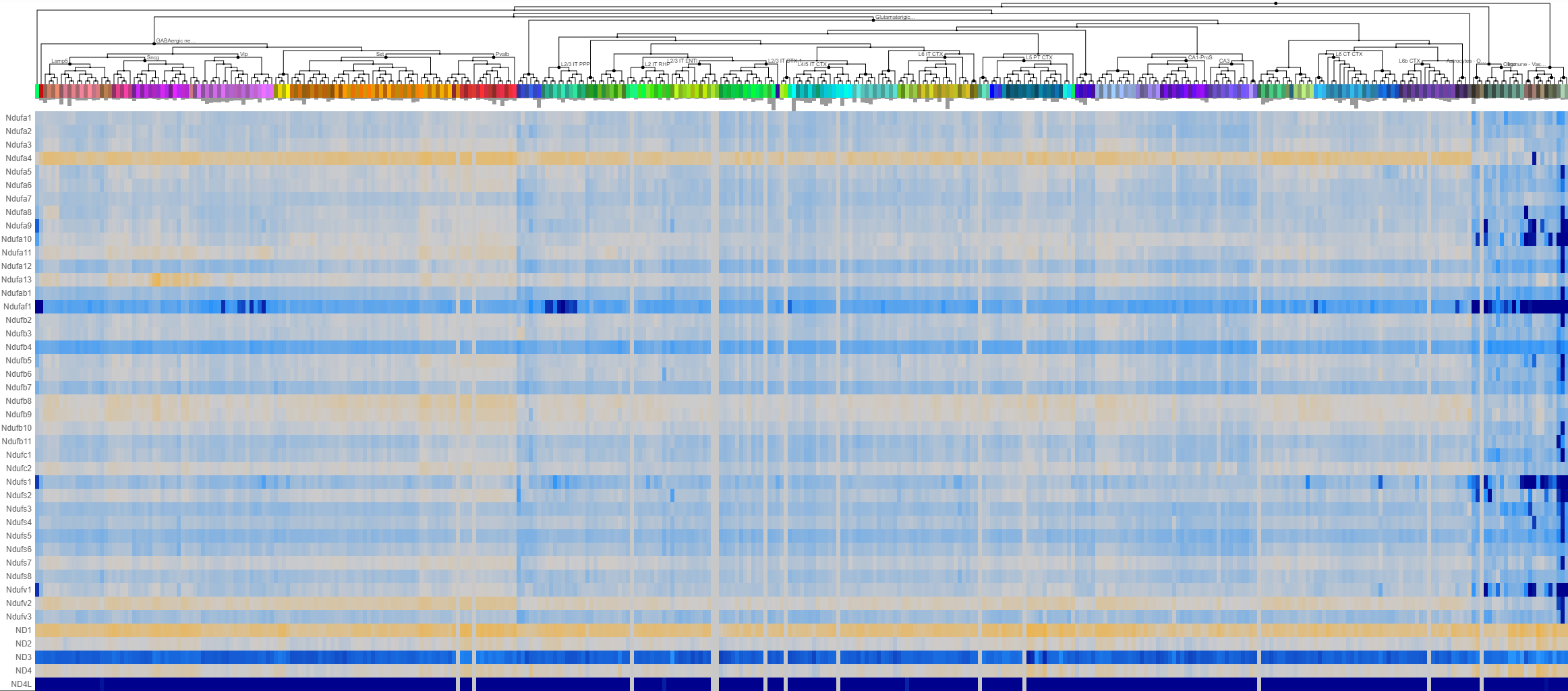

### mus complex II.png

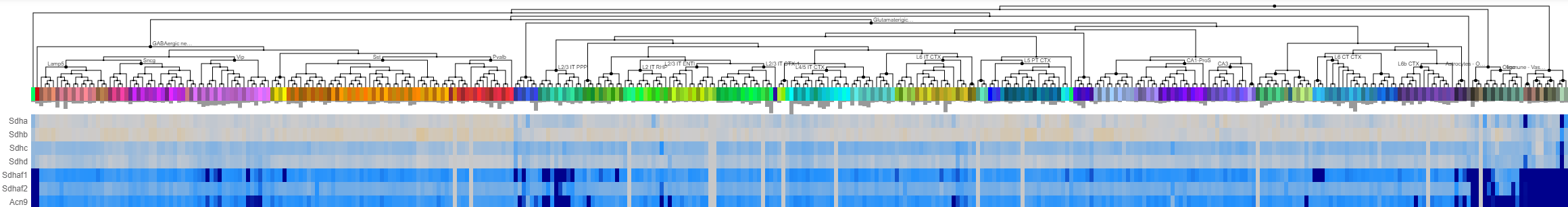

### mus complex III.png

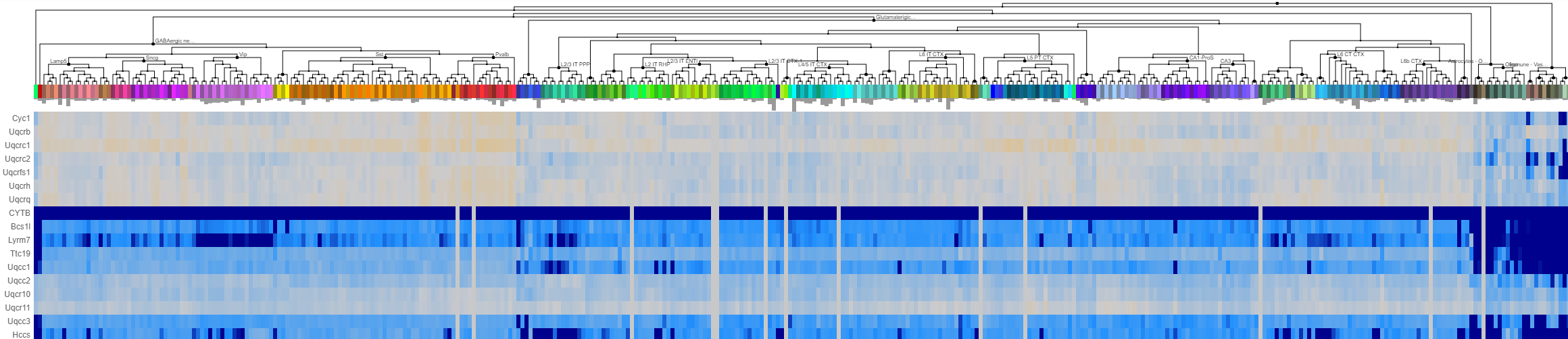

### mus complex IV.png

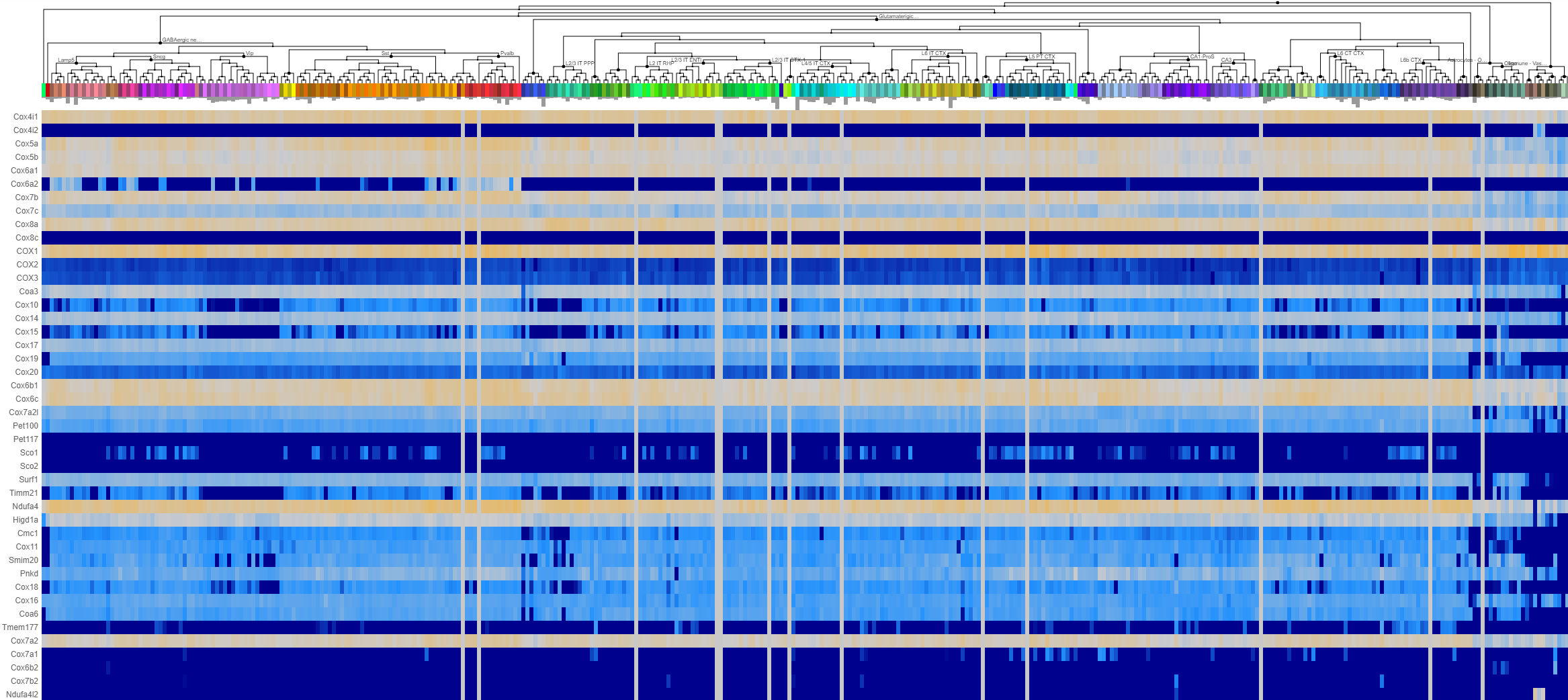

### mus complex V.png

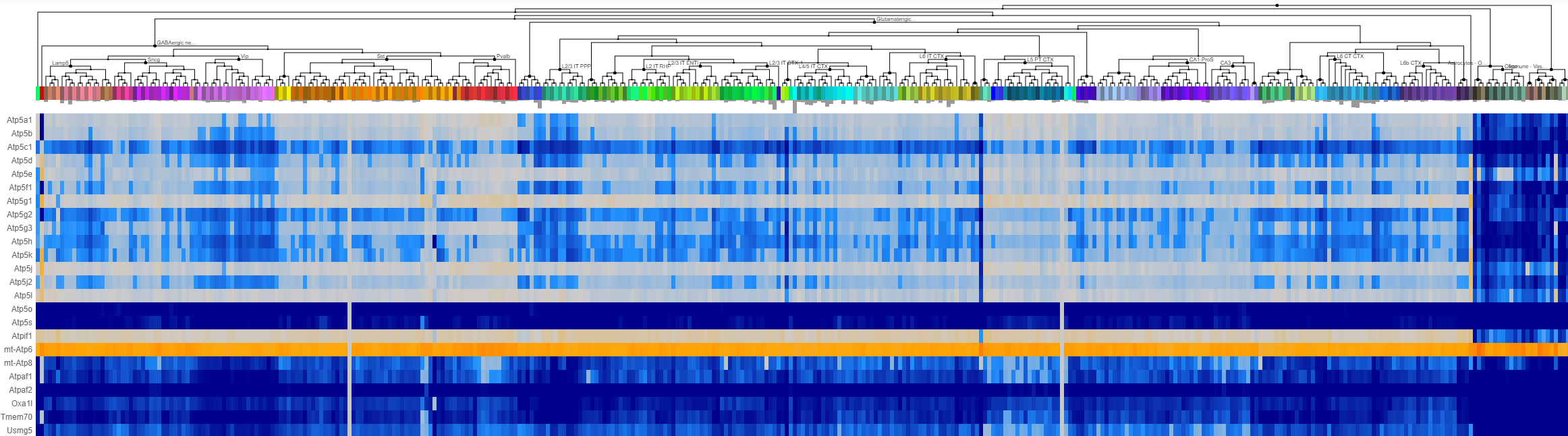

### mus cyt c.png

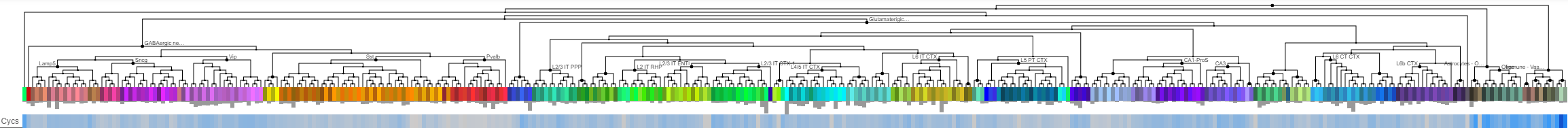

### mus MDH2 and IDH.png

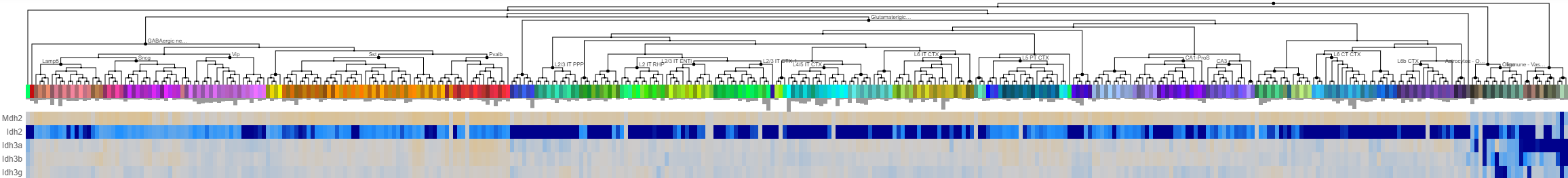

### mus PDH.png

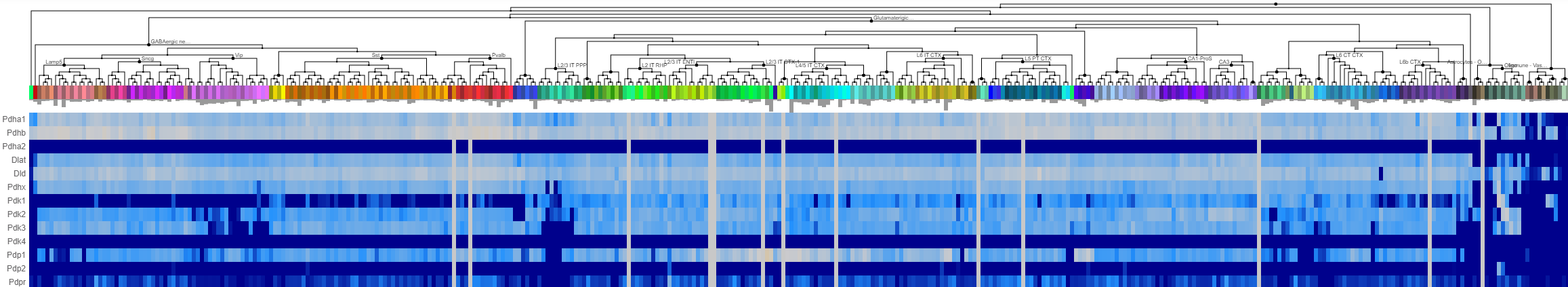

### mus Pi carriers and dicarboxylate carriers.png

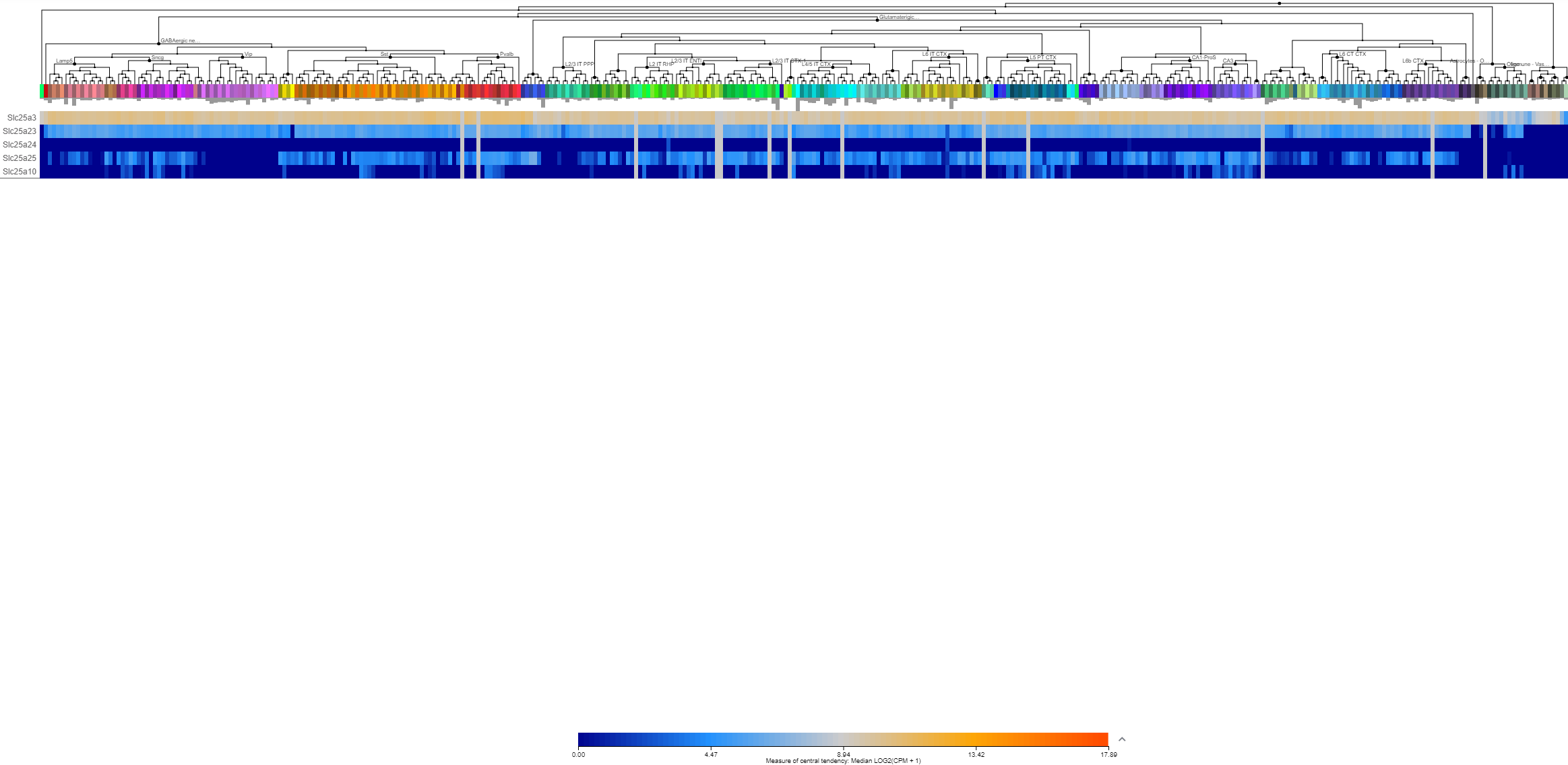

### Supplementary figure 1

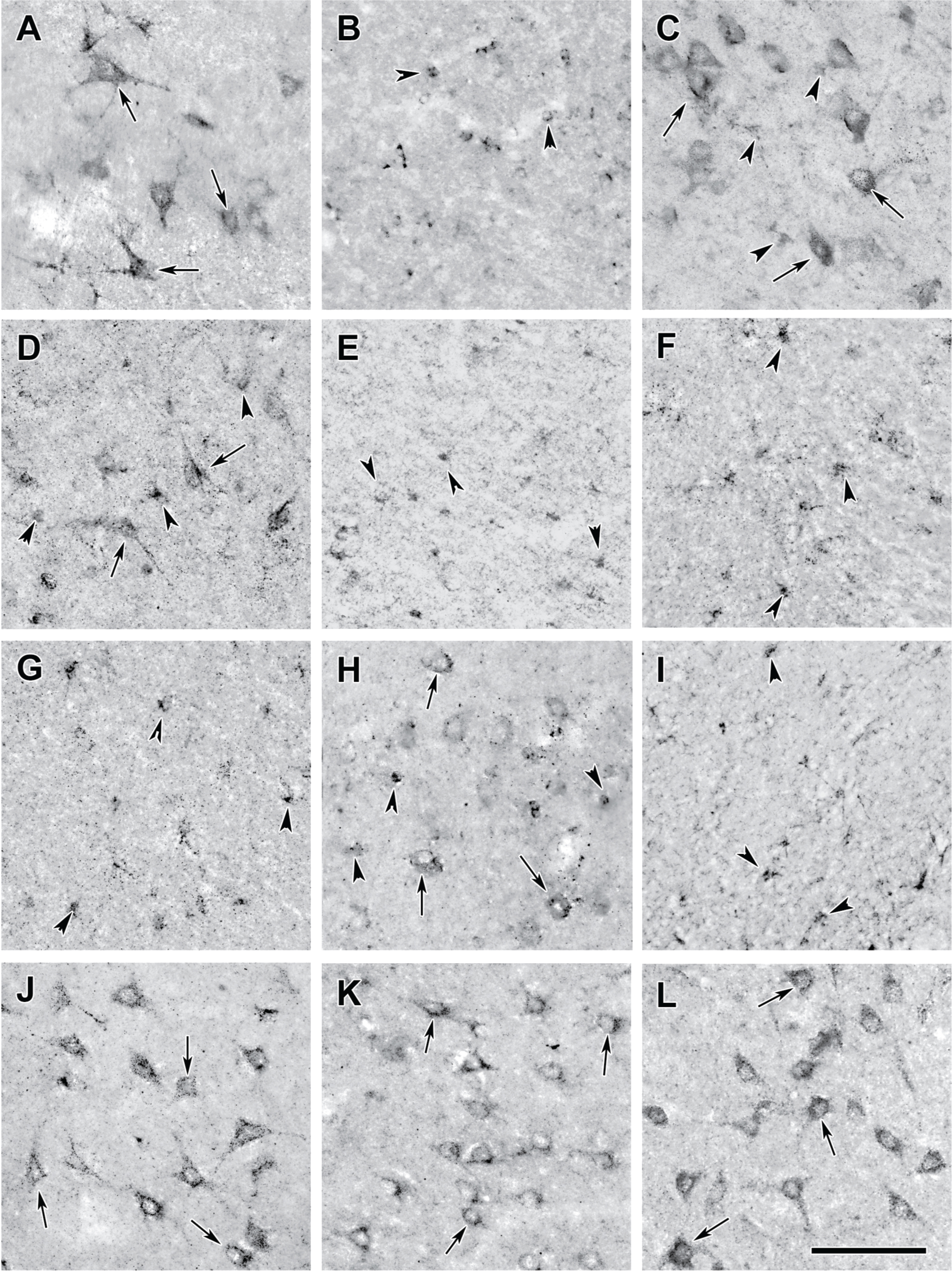

### Supplementary figure 2

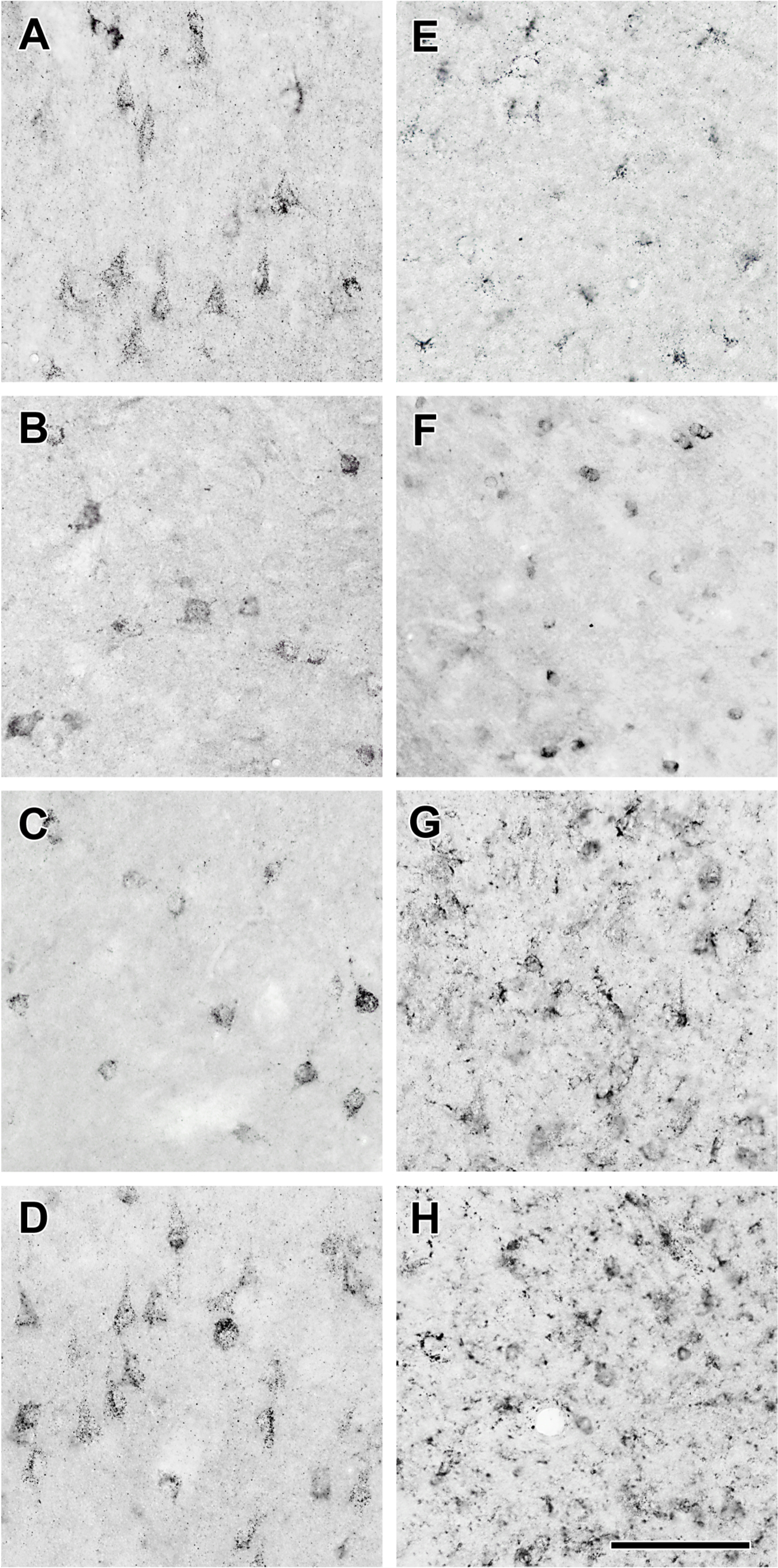

### Supplementary figure 3

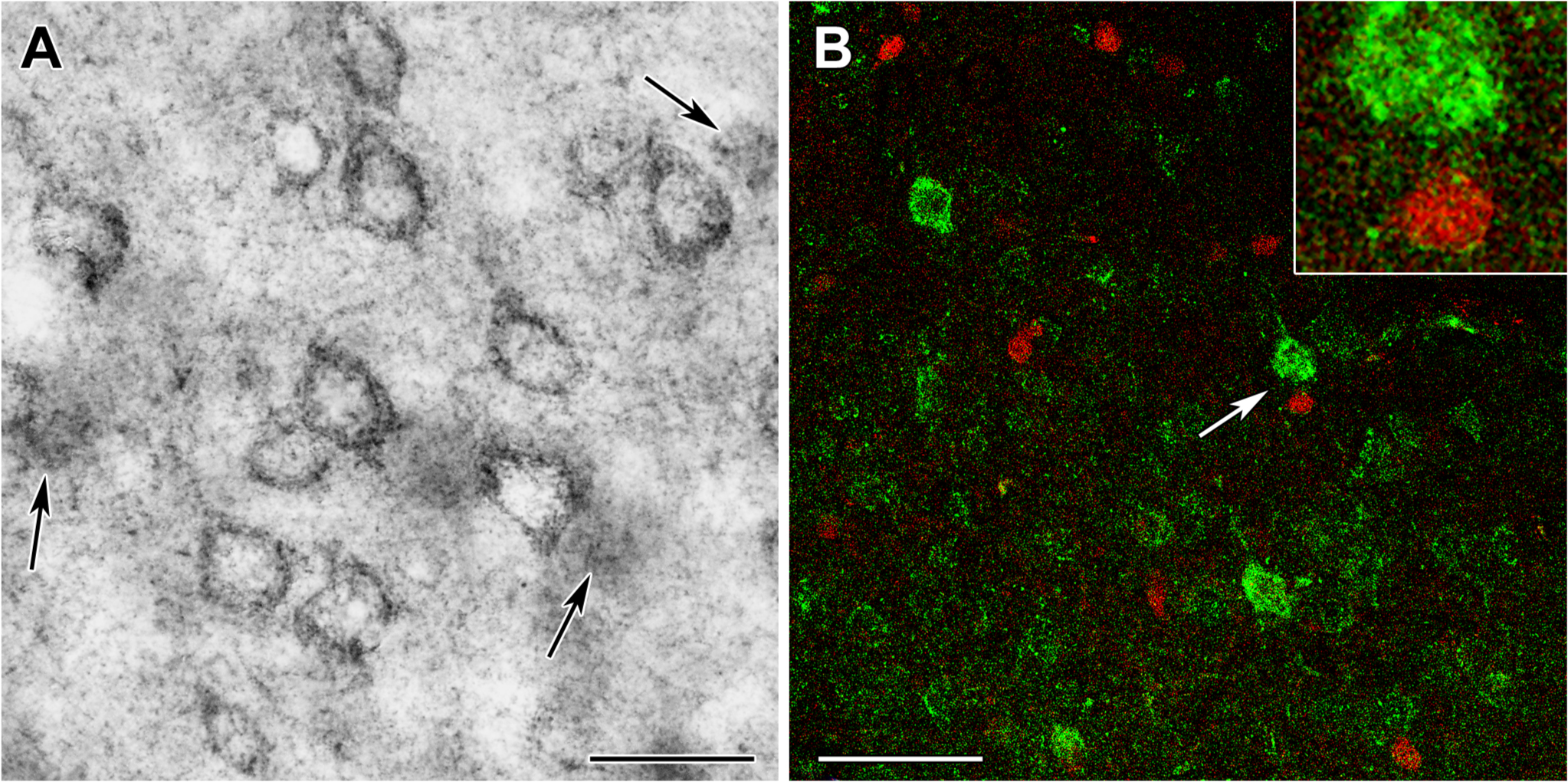
