## Supplementary table 1 for "Cell-specific expression of key mitochondrial enzymes precludes OXPHOS in astrocytes of the adult human neocortex and hippocampal formation"

| **Protein against which antibody was raised** | **Vendor**  **Cat no** | **Specific staining** | **RRID** | **Means of use**  **(DAB only or double labelling with S100)** | **Dilution** |
| --- | --- | --- | --- | --- | --- |
| PDH E2 | ab172617 | Yes | AB_2827534 | DAB only | 1:200 |
| PDH E1beta | ab155996 | Yes | AB_2814826 | Yes* | 1:500 |
| IDH2 | CST 56439 | Yes | AB_2799511 | DAB only | 1:100 |
| MDH2 | CST 11908 | Yes | AB_2797764 | DAB only | 1:100 |
| SDHA | ab137040 | No labelling | AB_2884996 |  |  |
| SDHB | ab175225 | Yes | AB_2904585 | DAB only | 1:500 |
| SDHC | ab155999 | Yes | AB_2810989 | DAB only | 1:500 |
| PDHX | ab155560 | Yes | AB_2924648 | DAB only | 1:100 |
| COX IV | ab14744 | No labelling | AB_301443 | - | 1:500 |
| COX IV | CST 4850 | Yes | AB_2085424 | Yes* | 1:500 |
| Cyt b-c1 complex subunit 9 | ab134909 | No labelling | AB_2924649 | - |  |
| Cytochrome C | CST 11940 | No labelling | AB_2637071 | - |  |
| Cytochrome c (6H2.B4) | CST 12963 | Yes | AB_2637072 | DAB only | 1:100 |
| Pyruvate carboxylase | 16588-1-AP | Yes | AB_1851513 | Yes* | 1:500 |
| MPC1 | CST 14462 | Yes | AB_2773729 | Yes* | 1:500 |
| MPC2 | CST 46141 | Yes | AB_2799295 | DAB only | 1:100 |
| alpha subunit ATPase | MS502 | No labelling | AB_478268 | - |  |
| S100B (D10G6) | CST 9550 | Yes | AB_10949319 |  | 1:1000 |

* yes: double fluorescent labelling with S100
